# Tracking *bla*_KPC_ Plasmid Dissemination within and between Enterobacterales across Michigan Over a Decade

**DOI:** 10.1101/2025.08.18.670900

**Authors:** Chaitra Shankar, Dhatri Badri Narayanan, Ashley Miihlbach, Sara McNamara, Brenda Brennan, Arianna Miles-Jay, Heather Blankenship, Auden Bahr, Evan S Snitkin

## Abstract

*bla*_KPC_ is endemic among Enterobacterales in the USA. While present on diverse plasmids, *bla*_KPC_ burden is often associated with the clonal spread of multi-drug resistant (MDR) epidemic lineages. In this study we sought to determine the relative contributions of clonal spread and plasmid transfer to *bla*_KPC_ burden across Michigan healthcare facilities over a decade. To this end we performed whole-genome sequencing of 1,058 KPC-producing isolates collected from 47 Michigan healthcare facilities between 2013 and 2022, including long-read sequencing for 527 isolates to enable precise plasmid tracking. Analysis with MOB-suite identified 64 distinct KPC plasmid types (“secondary clusters”), with the AK975 broad-host range plasmid being the most prevalent, found in 27% of isolates, spanning 20 species and 92 sequence types. Among genomes with AK975, 30% were from epidemic and 70% non-epidemic lineages, highlighting its broad role in regional *bla*_KPC_ spread. Epidemic lineages of various species constituted 46% of the study population. Epidemic lineages differed in their primary plasmids, and even within epidemic lineages there were clonal expansions with distinct *bla*_KPC_ plasmids, including in some cases AK975. These findings highlight two patterns of KPC spread: transmission of epidemic lineages harboring broad-range and lineage-specific KPC plasmids; and broader spread of AK975 among diverse species. Traditional surveillance studies often focus on common MDR lineages, potentially overlooking rare species and lineages that mediate the spread of plasmid-borne antimicrobial resistance (AMR) genes. Here we show how longitudinal studies tracking plasmids across species are essential to understand the pathways leading to AMR infections in hospitals.

**Importance:** This decade-long longitudinal study highlights the persistence and spread of key KPC-carrying plasmid across multiple bacterial species in the region, including some uncommon ones. It also emphasizes the differences in KPC plasmids across lineages within the same species. While some lineages acquire multiple plasmids with resistance, they are unable to successfully maintain the plasmids. In contrast, clonal sub-populations of KPC-producing bacteria disseminate selected plasmids, establishing a stable host-plasmid combination. Comprehensive genomic surveillance that includes all pathogenic species and plasmids is crucial to understanding the regional transmission dynamics of plasmid-borne antimicrobial resistance (AMR). While outbreak studies define the blowup of a successful lineage and associated plasmids, longitudinal studies identify the reservoir-species and circulating plasmids in the context of plasmid-borne AMR.

## Introduction

*Klebsiella pneumoniae* carbapenemases (KPC) have been the predominant carbapenemases in the United States since their initial identification in 2001, and they have been reported in multiple species of gram-negative bacteria (1). KPC production has been identified as an independent risk factor for mortality (2), with 28-day mortality rates as high as 39% (3). In 2017, the World Health Organization (WHO) identified KPC-producing *Enterobacterales* as high-priority pathogens, highlighting their association with limited treatment options and elevated mortality rates (4). To date, 245 variants of KPC have been identified among several species (https://www.ncbi.nlm.nih.gov/pathogens/refgene/#KPC), some with different susceptibilities to newer B-lactam/B-lactamase inhibitors such as ceftazidime-avibactam, meropenem-vaborbactam and imipenem-relebactam (5, 6).

Since its introduction, *bla*_KPC_ has spread throughout the world among multiple bacterial species with differences in the predominant lineage across diverse geographical regions. In the United States, the predominant KPC carrying lineages include *K. pneumoniae* ST258 and ST11 (7, 8) and *E. hormaechei* ST171 and ST78 (9, 10). These pathogens carry *bla*_KPC-like_ on the transposon Tn*4401*, and its variants, which are further mobilized by multiple plasmid types (11, 12). Prior to the COVID-19 pandemic, at least 259 plasmid types and three transposon families were linked to the mobilization of *bla*_KPC_ globally (13). The extensive genetic diversity associated with *bla*_KPC_ poses significant challenges for tracking and containing its spread.

Advances in long-read sequencing and hybrid genome assemblies have significantly enhanced the resolution of genomic studies, facilitating more accurate tracking of plasmid-borne and chromosomal antimicrobial resistance (AMR) gene transmission (14). However, monitoring the spread of AMR via plasmids remains challenging due to the complexity of plasmid biology, including the diversity of plasmid backbones, the presence of mobile genetic elements such as insertion sequences and transposons, and the capacity for plasmids to transfer across multiple bacterial species (2). To address these challenges, surveillance strategies should incorporate longitudinal sampling across diverse bacterial species to better understand the establishment and dissemination of AMR plasmids within a region. However, most studies investigating the spread and dynamics of *bla*_KPC_ are limited in scope; often constrained by short study periods, a focus on a single species, and/or data collected from a single healthcare facility. Even in instances where multiple species are included, the analyses frequently remain confined to a single institution or few institutions (3–5). These limitations hinder a comprehensive understanding of how diverse species and lineages influence plasmid dissemination within a broader regional context.

The present study addresses several of these gaps by employing a longitudinal approach spanning ten years, encompassing 47 healthcare facilities across the state of Michigan, and including all *bla*_KPC_-positive species identified during this period. Through the generation of more than 500 hybrid genome assemblies, and tracking KPC over time, we observed extensive dynamics in lineage prevalence over time, the often transient associations of lineages with plasmids and the central role played by a single broad-host range plasmid in spreading KPC among diverse Enterobacterales species over space and time.

## Results

### Overview of the study data

Among the 1,065 carbapenemase-producing bacteria analyzed in this study, four major species/species complexes—*Escherichia coli*, *Klebsiella pneumoniae* species complex, *Enterobacter cloacae* complex, and *Citrobacter freundii* complex - accounted for 91% of the total isolates. The prevalence of isolates and the major epidemic lineages during the study period are shown in **Fig 1**. Among the 1,065 genomes that were included after quality control, 7 genomes lacked *bla*_KPC_ and hence were excluded from the study. Chromosomal *bla*_KPC_ alleles were observed among 8% (n=90) of the isolates (**Suppl. Fig 1**) among which 16 genomes carried multiple copies of *bla*_KPC_ on both the chromosome and plasmids. There were five variants of *bla*_KPC_ observed, among which *bla*_KPC-2_ (70%, n=786) was the most predominant, followed by *bla*_KPC-3_ (26% , n=297) (**Suppl. Fig 1**). Further, *bla*_KPC-2_ was carried on multiple Tn*4401* variants among which Tn*4401a* (28%, n=220) and Tn*4401b* (70%, n=548) were common. *bla*_KPC-3_ was predominantly carried on Tn*4401b* (97%, n =287) and Tn*4401d* (3%, n = 10) (**Fig 1b**).

**Figure 1:**
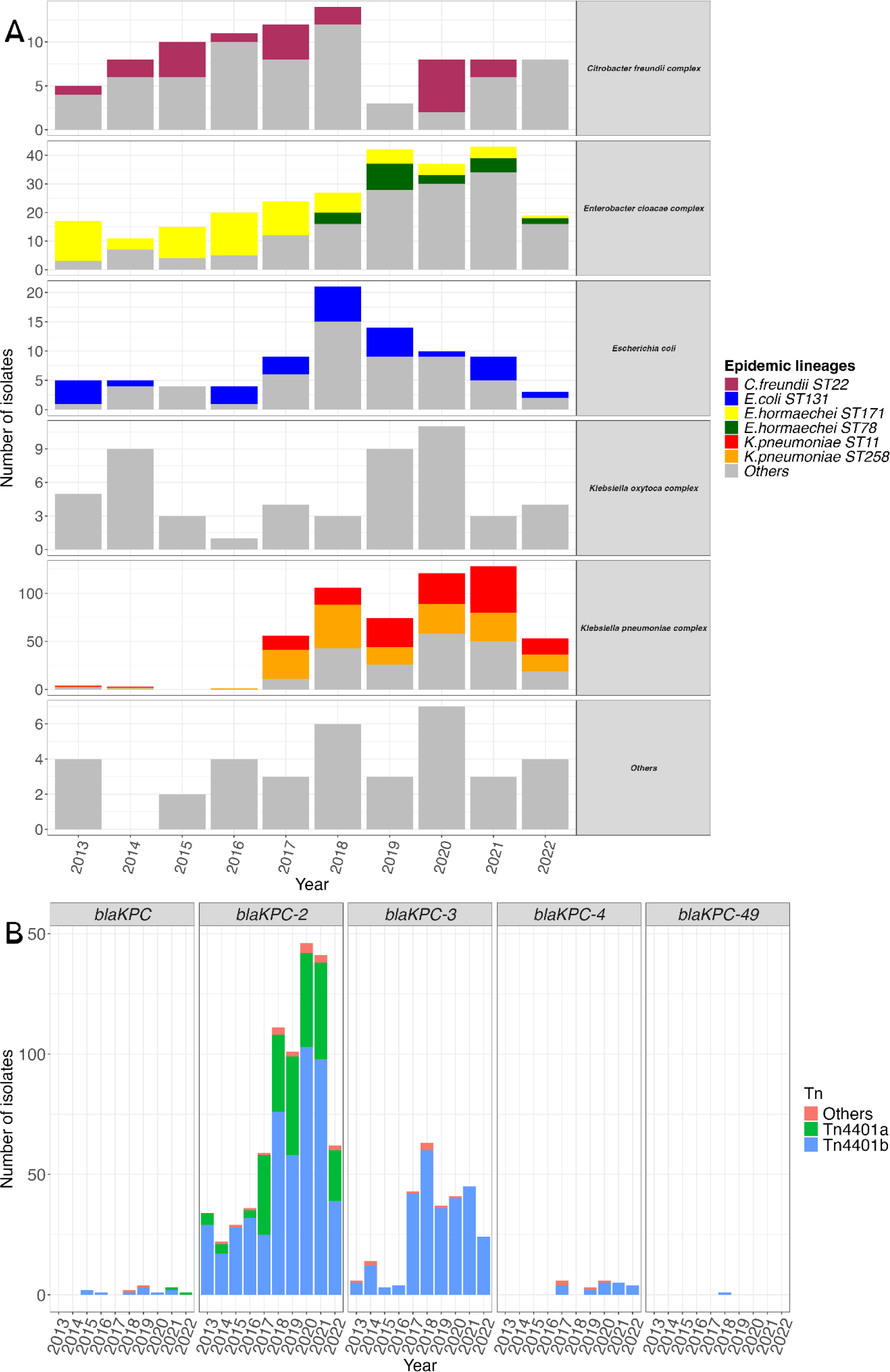
Overview of species, STs and *bla*_KPC_ alleles. **A)** Distribution of KPC producing bacteria and the associated epidemic lineages during the study period is shown. Six epidemic lineages among the *bla*_KPC_ carrying species were identified, which comprised 46% of the study isolates. While most epidemic lineages were present throughout the study period in varying proportions, a clonal shift was observed among *E. hormaechei*. *E. hormaechei* ST171 declined 2018 onwards which was accompanied by increase in *E. hormaechei* ST78. **B)** Association of *bla*_KPC_ alleles and transposons is shown over time. *bla*_KPC-2_ and *bla*_KPC-3_ were the predominant alleles in the study isolates. While *bla*_KPC-2_ was mobilized by both Tn*4401a* (28%) and Tn*4401b* (70%), 97% of *bla*_KPC-3_ was mobilized by Tn*4401b*. ST11: *K. pneumoniae* ST11, ST131: *E. coli* ST131, ST171: *E. hormaechei* ST171, ST22: *C. freundii* ST22, ST258: *K. pneumoniae* ST258 and ST78: *E. hormaechei* ST78.

**Figure 2:**
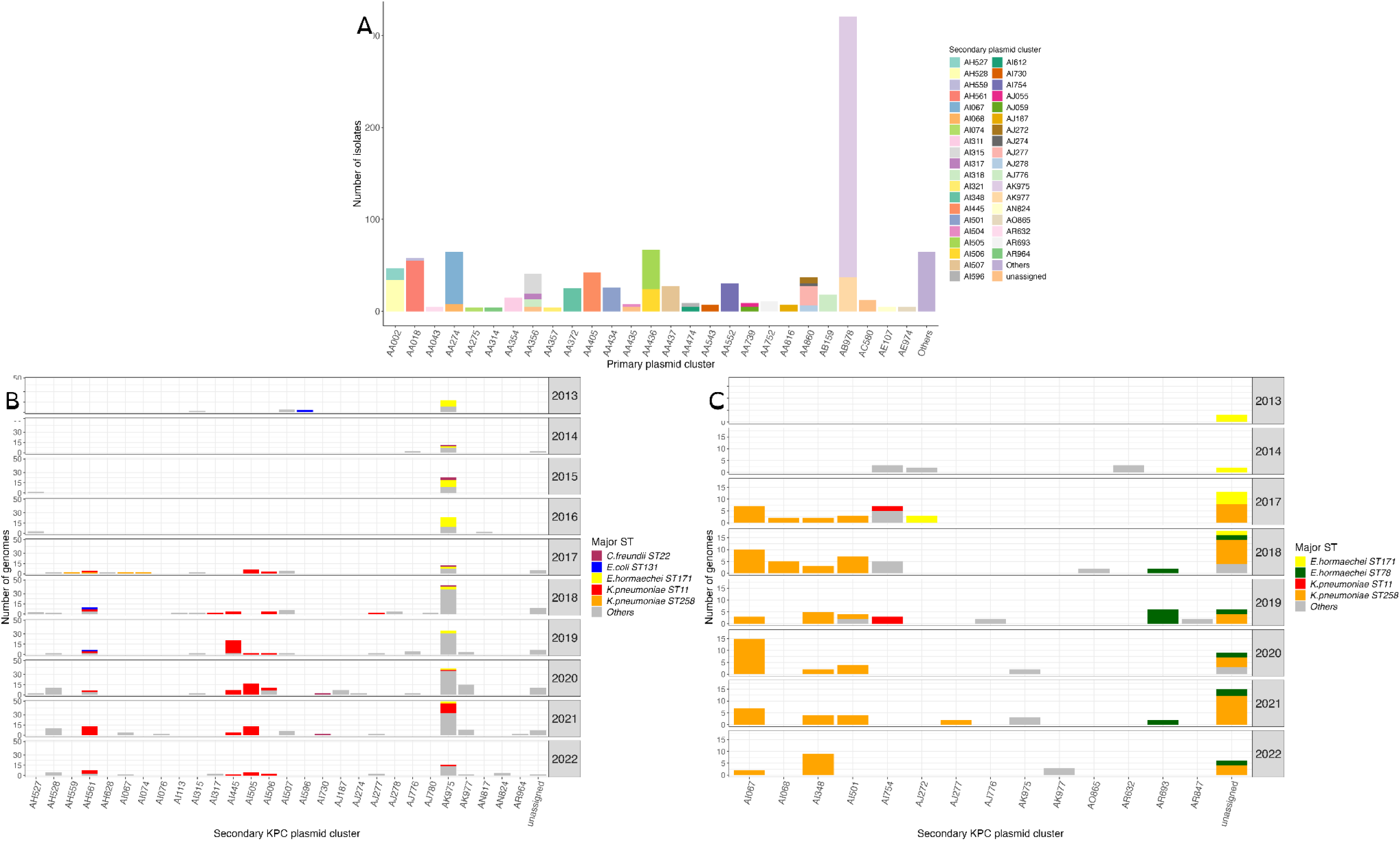
Classification and distribution of KPC plasmids in the study. **A)** Primary and secondary KPC plasmid clusters for study population as obtained from MOB-suite are shown. Plasmid clusters with <2 plasmids were grouped as others. A single primary plasmid cluster is classified into one or more secondary clusters. AB978 was the key primary plasmid cluster associated with *bla*_KPC_ which comprised two secondary clusters AK975 and AK977. **B)** The common secondary plasmid clusters associated with *bla*_KPC-2_ are shown. Notably, AK975, the dominant KPC cluster, was present among epidemic as well as non-epidemic lineages. Plasmid clusters that occurred just once during each year among a single epidemic lineage were excluded from the figure. *E. hormaechei* ST78 lacked *bla*_KPC-2_. Prevalence of AH561 plasmid, increased among *K. pneumoniae* ST11 from 2019 onwards and its presence drastically reduced among other lineages. **C)** The common secondary plasmid clusters associated with *bla*_KPC-3_ are shown. *K. pneumoniae* ST258 and *E. hormaechei* ST78 were the chief epidemic lineages associated with *bla*_KPC-3_. These two lineages carry distinct plasmid clusters mobilizing *bla*_KPC-3_. Plasmid clusters that occurred just once during each year among a single epidemic lineage were excluded from the figure.

Long-read sequencing data was generated using Oxford Nanopore Technology (ONT) for 599 genomes, selected to include representatives of all species, ST, *bla*_KPC_ allele and plasmid type combinations from each year of the study. Among these 599 genomes, 527 had resultant hybrid assemblies that passed quality control and were used to update the MOB-suite database to enhance predictions of locally circulating plasmids. After addition of hybrid plasmids to the database, 71% (n=1840) of plasmids matched to a hybrid plasmid as the reference as shown in **Suppl. Fig 2**. There was also a significant shift in mash distances between predictive plasmids and their best match reference in the database. The mean and the inter-quartile range (IQR) for the initial MOB-suite run was 0.088 and 0.071, respectively; while for the updated run, the mean was 0.017 and IQR was 0.023. The shift in mash distance (*p* < 0.005 for mean mash distance), demonstrates that by updating the reference database with locally circulating plasmids we significantly improved our ability to map short-read assemblies to matching plasmids (**Suppl. Fig 2**).

After running MOB-suite, a total of 974 KPC plasmids were identified from the study isolates, which included 393 complete circular plasmids obtained through hybrid assembly. The *bla*_KPC_ carrying plasmids were diverse, with 69 primary clusters and 64 secondary clusters according to MOB-suite classification and are shown in **Fig 2a**. The most predominant secondary plasmid clusters associated with *bla*_KPC-2_ and *bla*_KPC-3_ among the various species are shown in **Fig 2b and Fig2c** and the primary clusters are shown in **Suppl. Fig 2**. The most common secondary plasmid cluster associated with *bla*_KPC-2_ (n=741) was AK975 (n=275, 37%). Among the 280 *bla*_KPC-3_ carrying isolates, the most common secondary plasmid cluster was AI067 (n=46, 16%). The corresponding primary clusters classified by MOB-suite as shown in **Suppl. Fig 3.** Incompatibility types and results for the KPC plasmids of the present study are mentioned in **Suppl. Fig 4**. The plasmid AK975, corresponded to rep_cluster_1367 (n=291, 30%). The second most common incompatibility type was the IncF group with rep_cluster_2183(n=163, 17%).

### Contribution of epidemic lineages in spread of *bla*_KPC_ in Michigan

#### *Klebsiella pneumoniae* species complex *(KpSC)*

Among the 546 KpSC isolates, a total of 75 sequence types (STs) were identified. The two predominant epidemic lineages associated with KPC were *K. pneumoniae* ST258 (31.6%, n = 174) carrying *bla*_KPC-3_, and *K. pneumoniae* ST11 (29.4%, n = 162) carrying both *bla*_KPC-2_ and *bla*_KPC-3_. There were notable differences in the KPC plasmid clusters between these two epidemic lineages. In *K. pneumoniae* ST11, the most prevalent KPC plasmid cluster was AI505 (27.7%, n = 45), followed by AI445 (24%, n = 39) and AH561 (16.6%, n = 27). AI505 was exclusive to *K. pneumoniae* ST11 while AI445 and AH561 were infrequently present among other lineages of *K. pneumoniae* and *E. coli*. In contrast, *K. pneumoniae* ST258 was primarily associated with AI067 (28%, n = 49), followed by AI348 (15.5%, n = 27). AI067 was seen in one other non-ST258 isolate while AI348 was observed in 7 non-ST258 isolates. Nearly a quarter of the *K. pneumoniae* ST258 were predicted to carry *bla*_KPC-3_ on chromosomes (24%, n = 41) which are represented as unclassified secondary clusters in **Fig 3a**. Among KpSC, AK975, the most prevalent KPC plasmid cluster in this study, was detected among 12% (n=66/546) of isolates across multiple STs, including a small proportion of *K. pneumoniae* ST11 isolates (n=19/162, 12%). Hence among KpSC, *K. pneumoniae* ST11 showed evidence of plasmid sharing with other species and lineages, while *K. pneumoniae* ST258 was conserved with specific plasmid clusters. In addition, the plasmids frequently found with epidemic lineages were different from the ones that were carried by non-epidemic lineages.

**Figure 3:**
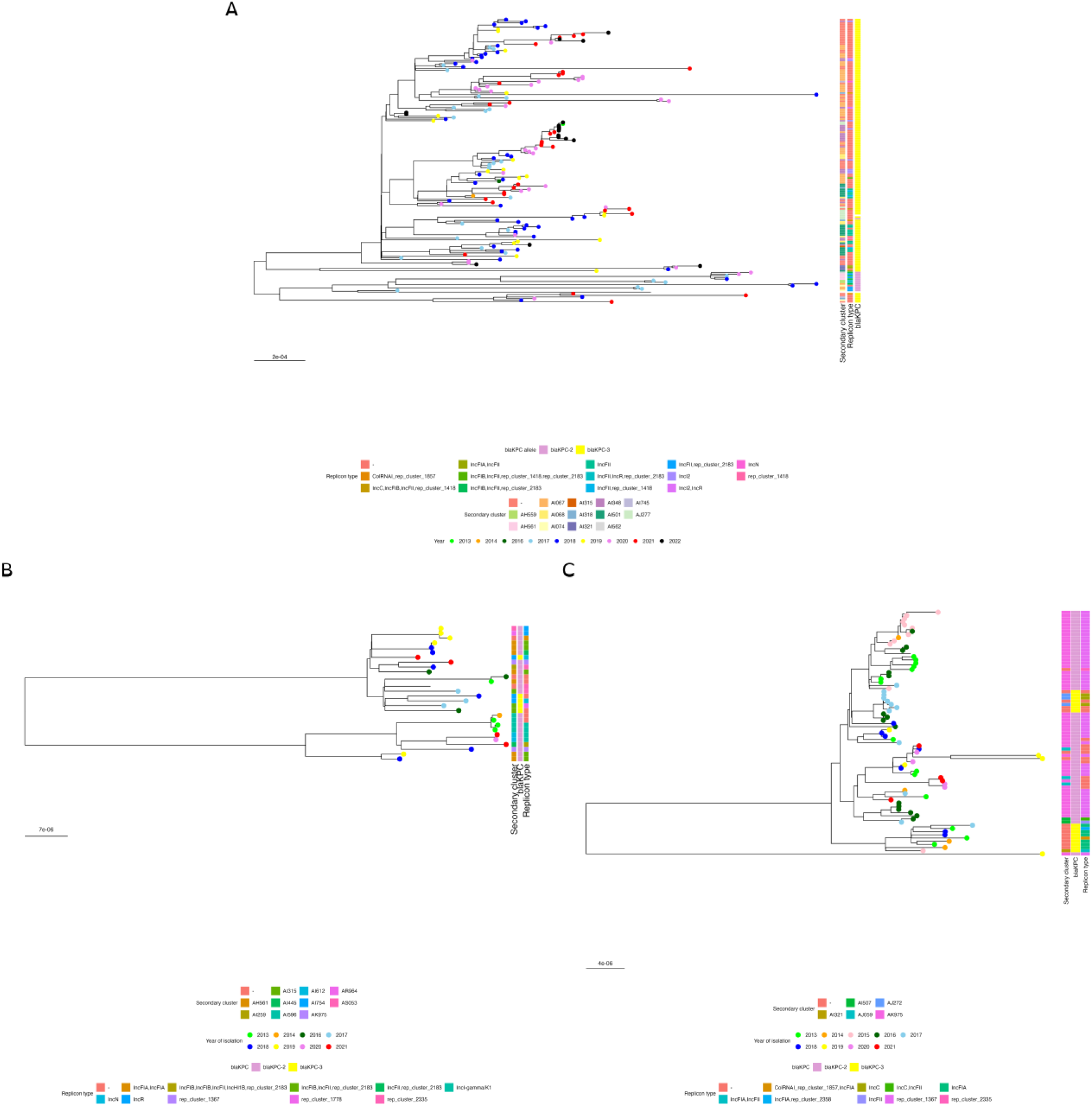
Core genome phylogeny of epidemic lineages of KPC carrying species. **A)** Core genome phylogeny of *K. pneumoniae* ST258 (n=174). *K. pneumoniae* ST258 almost exclusively harbors *bla*_KPC-3_. The diversity of KPC-carrying plasmids within this lineage reflects its genetic plasticity in acquiring multiple plasmids. However, there was a lack of predominant KPC-plasmid that was widely transmitted. The most common KPC-plasmid within the lineage, AI067, accounted for only 28%. A small set of *bla*_KPC-2_ carrying isolates were observed that were distantly related to the *bla*_KPC-3_ carrying isolates. **B)** Core genome phylogeny of *Escherichia coli* ST131 (n=28). Two main clades were observed in *E. coli* ST131. There is a significant plasmid diversity observed, with key KPC-plasmid, AH561, accounting to only 25%. Ten KPC-plasmid clusters were identified in this lineage. **C)** Core genome phylogeny of *E. hormaechei* ST171 (n=77). *E. hormaechei* ST171 shows 3 main clades with two carrying *bla*_KPC-2_ and the 3rd, relatively distinct clade, carrying *bla*_KPC-3_. When compared to other epidemic lineages, ST171 is conserved with KPC-plasmids. AK975,the predominant KPC-plasmid identified in the study, accounts to 71% of KPC-plasmid in this lineage.

#### *Enterobacter cloacae* complex *(ECC)*

Among the 255 ECC isolates, a total of 50 diverse sequence types (STs) were identified, with *E. hormaechei* ST171 (30%, n = 77) and *E. hormaechei* ST78 (9%, n = 23) emerging as the predominant lineages. *E. hormaechei* ST171 was dominant between 2013 and 2016, during which ST78 was absent. However, ST78 emerged in 2018 and subsequently reached proportions comparable to ST171. The two lineages differed in their *bla*_KPC_ alleles and associated plasmids. While 80% (n = 60) of ST171 isolates encoded *bla*_KPC-2_, 96% (n = 22) of ST78 isolates encoded *bla*_KPC-3_. AK975 was the predominant KPC plasmid cluster in ECC, observed in 71% of ST171 isolates and 47% of non-epidemic lineages (**Fig 3c**). Of note is that ST171 was an early carrier of AK975 in the region, indicating its potential role as an early reservoir. In contrast to ST171, *E. hormaechei* ST78 lacked AK975 and carried 52% of the *bla*_KPC-3_ on AR683, a plasmid exclusive to this lineage.

#### *Citrobacter freundii* complex (CFC)

Among the 87 CFC isolates, 34 sequence types (STs) were identified, with *C. freundii* ST22, the predominant epidemic carbapenemase lineage, accounting for 25% (n = 22) of the isolates. The *bla*_KPC-2_ allele was highly prevalent in this group, comprising 81.6% (n = 71) of all CFC isolates. A diverse range of KPC-associated KPC plasmid clusters was observed among CFC. However, AK975, the endemic KPC plasmid, was carried by 34% (n=29) of CFC including half of *C. freundii* ST22 (n=12/22, 54.5%).

#### Escherichia coli

Eighty-four *E. coli* isolates were present in the study collection and the epidemic lineage, *E. coli* ST131, comprised 33.3% (n=28). Twenty-eight diverse sequence types were observed carrying *bla*_KPC-2_ (75%, n=63) and *bla*_KPC-3_ (23.8%, n = 20) with a single isolate carrying *bla*_KPC-4_. There were multiple plasmid clusters associated with *bla*_KPC_ alleles and AK975 was seen among 21% (n=18). *E. coli* ST131 exhibited significant diversity in KPC-plasmid associations, with multiple plasmid clusters identified (**Fig 3b**). Notably, the most prevalent cluster, AH561, accounted for only 25% (n = 7) of ST131 isolates, highlighting the absence of a dominant plasmid type and the broad variability in KPC-plasmid distribution within this lineage.

### Determination of clonal spread among the epidemic lineages

Since the epidemic lineages among the major species constituted 46% (486/1058) of the study population, determining the role of these lineages in amplifying specific KPC plasmids was essential. In particular, we sought to identify the proportion of isolates belonging to epidemic lineages that we could link to local transmission and determine how epidemic lineages contributed to the spread of KPC plasmids. To this end, thresholds for defining recent transmission were established for each lineage based on the distribution of single-nucleotide variant (SNV) distances of intra- and inter- healthcare facility isolate pairs (**Suppl. Fig 5**). For *E. coli* ST131, *K. pneumoniae* ST11 and *K. pneumoniae* ST258, the threshold was set to 6 SNPs, for *C. freundii* it was set to 4 SNPs, and for *E. hormaechei* ST78 and *E. hormaechei* ST171 it was set to 5 SNPs. Overall, among the 486 isolates belonging to the aforementioned epidemic lineages, 64% (n=311) were linked to another isolate below the established threshold.

We also noted that within epidemic lineages, there were clonal expansions associated with different KPC plasmid clusters (**Fig 3 and Suppl. Fig 6)**. In particular, among each of the epidemic lineages, there were multiple sub-lineages, with some of these being restricted in time (isolated during a single year among multiple healthcare facilities). These sub-lineages exhibited both intra- and inter-healthcare facility transmission, as illustrated in the pairwise SNP distance matrices (**Suppl. Fig 5**). Among *Klebsiella pneumoniae* ST11 and ST258, several sub-lineages were associated with specific KPC plasmids. In contrast, *Escherichia coli* ST131 harbored multiple KPC plasmid types, none of which were successfully disseminated. In *E. hormaechei* ST171, multiple sub- lineages effectively spread the predominant KPC plasmid, whereas in *E. hormaechei* ST78, a variety of plasmids were associated with different sub-lineages. Furthermore, clades of *K. pneumoniae* ST11, *K. pneumoniae* ST258 and *E. hormaechei* ST171 carrying non-predominant *bla*_KPC_ alleles were phylogenetically distinct from other sub- lineages. *C. freundii* ST22 had sub-lineages associated with the spread of predominant KPC plasmid.

### Inter-species KPC plasmid sharing

Hybrid assemblies yielded 393 complete circular KPC plasmid sequences. A mash distance-based phylogeny, constructed using Mashtree (**Fig 4a**), revealed that while the major KPC plasmid cluster was AK975, additional clades contained other plasmid types with low mash distance, several detected in later study years between 2020 and 2022.

**Figure 4:**
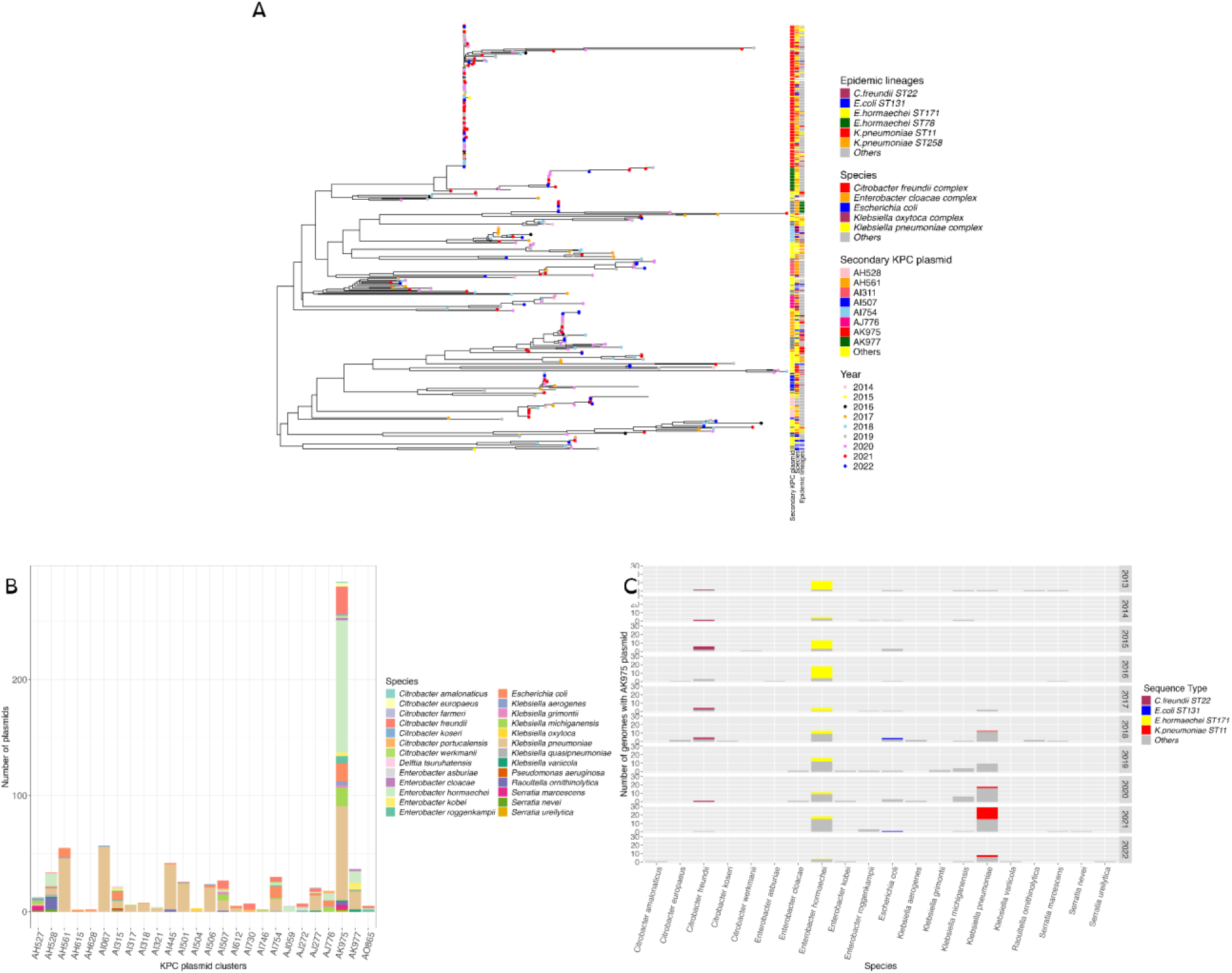
Distribution of KPC plasmids among species and lineages. **A)** A phylogeny of all complete circular KPC plasmids was constructed using Mashtree (n=393). Secondary KPC plasmid refers to the secondary plasmid clusters defined by MOB-suite. Apart from the most common clusters, remaining clusters were grouped together as “Others”. Most KPC plasmids are present among multiple species and frequently shared among epidemic and non-epidemic lineages. **B)** Among a total of 64 secondary plasmid clusters identified by MOB-suite to carry *bla*_KPC_, 26 plasmid clusters were present in >1 species. Plasmids present in single species were excluded from this figure. On average, each of the plasmids in the figure were present in 3 species. **C)** Distribution of AK975 KPC plasmid during the study among bacterial species and sequence types. Others: Non-epidemic lineages of all species.

The most common KPC plasmid types were shared across species, with few plasmid types restricted to specific species or lineages as shown in **Fig 4b**. Notably, **Fig 4 and Suppl. Fig 7**, highlight the role of non-epidemic lineages and uncommon species in plasmid dissemination. For example, the plasmid, AK975 was found across both epidemic and non-epidemic lineages, as well as in uncommon pathogenic species including *Citrobacter werkmanii*, *Citrobacter europaeus*, and *Klebsiella grimontii*. More broadly, 26 distinct KPC plasmid clusters **(Fig 4b)** were found to be carried by multiple species (range: 2-20 species, median: 3 species and inter-quartile range: 3 species), emphasizing the broad host range and mobility of these plasmids. In addition, we noted that the KPC plasmids carried additional antimicrobial resistance genes, which are shown in **Supl. Fig 8**. Overall, this highlights the importance of multi-species surveillance for plasmid-borne antimicrobial resistance.

Further analysis of the major KPC secondary cluster, AK975 was performed using SNP-based comparisons against its closest reference plasmid from the MOB-suite database. The resulting phylogeny of 130 AK975 KPC-carrying plasmids (**Suppl. Fig 9**) showed a lack of genetic variation, with 51% of the pairwise comparison of AK975 differing by 0 SNPs. While the small size of this plasmid (median 43.6 Kb) makes the lack of genetic variation expected, it demonstrates how genetic changes on the plasmid do not appear necessary to reside in different genetic backgrounds. **Fig 4c** illustrates the broad distribution of this plasmid, which was identified in 20 species and 92 diverse sequence types. *E. hormaechei* emerged as a consistent host throughout the study, contributing significantly to plasmid dissemination. Diverse *K. pneumoniae* lineages were primarily involved during the second-half of the study period. In addition to the clonal expansion of epidemic lineages, there is evidence of regional spread and persistence of an endemic KPC plasmid.

## Discussion

The present study provides insights into the prevalence of KPC producing species, lineages and plasmids that were observed across Michigan healthcare facilities from 2013 to 2022. This longitudinal study helps to address knowledge gaps in regional genomic epidemiology by describing the patterns of KPC spread and transmission in the state. We observed two phenomena driving the spread of KPC in the region: (i) epidemic lineages, often with sub-populations harboring distinct KPC plasmids, clonally disseminating across the region and (ii) a single broad-host range plasmid, AK975, spreading among diverse species and lineages, with varying degree of clonal spread following plasmid acquisition. Together, these results highlight the interplay between epidemic lineages and widespread sharing of AK975, a multi-species regional plasmid, in dissemination and maintaining *bla*_KPC_ in Michigan.

Key global lineages associated with *bla*_KPC_, such as *K. pneumoniae* ST258, *K. pneumoniae* ST11, *E. hormaechei* ST171, and *E. coli* ST131, were observed in our study, representing 46% of the total population (8–11). Notably, despite their stable presence in aggregate, there was variation over time with respect to which lineages dominated, and the KPC-plasmids they carried. For example, a significant shift was observed within *E. hormaechei* from ST171 to ST78 lineage starting in 2017, indicating a clonal change within the species. This shift in turn was accompanied by changes in the KPC allele and the associated plasmid types (**Fig 3c and Suppl Fig 5b**). Moreover, even within stably present lineages like *K. pneumoniae* ST258, we identified small clonal populations linked to specific plasmid clusters, highlighting variation over time in the plasmid on which its defining resistance gene is carried. Exhibiting even more variation was *E. coli* ST131, which showed evidence for multiple independent acquisitions of diverse KPC plasmids, with limited onward spread. These findings highlight the dynamic nature of carriers mediating the spread of antibiotic resistance, emphasizing the growing challenge of surveilling for ongoing outbreaks and emerging threats.

Genomic surveillance can inform the origins and molecular basis for plasmid-borne antimicrobial resistance genes (ARGs) in a region, discriminating between clonal spread of a single epidemic lineage, sudden introduction and spread of a sporadic lineage, or plasmid spread among multiple species of bacteria (11, 13). In the present study, a combination of clonal spread of lineages and spread of endemic plasmid among multiple lineages contributed to the spread of *bla*_KPC_. The endemic KPC plasmid, AK975, was shared among multiple epidemic lineages including *K. pneumoniae* ST11, *E. coli* ST131, *E. hormaechei* ST171, as well as numerous non-epidemic lineages. In parallel to this sustained regional plasmid outbreak, *K. pneumoniae* ST258, which was never found associated with this major plasmid, propagated KPC via the AI067 plasmid, which was almost exclusive to this lineage. The KPC plasmid AI067 was only found in eight other genomes, with little to no onward spread following acquisition of this plasmid. Similarly, *E. hormaechei* ST78 harbored the KPC plasmid AR683, which was exclusive to this lineage. A similar longitudinal study from New York, reports the clonal dissemination of this KPC plasmid among *E. hormaechei* ST171 and *K. pneumoniae* ST258 (15). In the New York study, *K. pneumoniae* ST258 was the most diverse lineage carrying up to 18 KPC plasmid types, which is comparable to our study, wherein this lineage carried 14 distinct KPC plasmids (15). Chromosomal integration of *bla*_KPC_ in *K. pneumoniae* ST258 was also reported, consistent with our observations. Another longitudinal study from Canada exploring the epidemiology of *bla*_KPC_ producing Enterobacterales also reports the epidemic lineages *K. pneumoniae* ST11, *E. coli* ST131 and *C. freundii* ST22 and long-term persistence of 3 KPC plasmids (16). Similar to our observations of multiple pathways of *bla*_KPC_ spread, Lerminiaux and colleagues report a combination of clonal spread as well as horizontal transmission of KPC plasmids (16).

AK975, the major endemic KPC plasmid, was conserved in its size across species, with 95% of the complete plasmids sized 43Kb, and co-carried *bla*_TEM-1_. Similarly, two predominant KPC plasmid clusters in the Canadian study were conserved in size compared to others in the study and carried other ARGs in addition to *bla*_KPC_ (16). The enduring success of a KPC-plasmid over a decade within a regional ecosystem likely results from a confluence of genetic and epidemiological factors. KPC plasmids like AK975, frequently exhibit broad host ranges and possess efficient transfer and maintenance mechanisms, facilitating dissemination across diverse Enterobacterales species in both healthcare and community settings(11, 13, 15). Notably, their carriage tends to incur minimal fitness costs to bacterial hosts, supporting long-term persistence even in the absence of antibiotics (17, 18). Since AK975 is <50Kb and carries two ARGs, it likely does not impose as high a fitness cost on the host as compared to other KPC plasmids that can be up to 1.2Mb (19). In addition, since it carries two ARGs, the deletion of these may not significantly impact the fitness-cost, potentially contributing to the stable association of this plasmid with these important AMR determinants (20–22). This plasmid has demonstrated remarkable genetic stability across diverse bacterial hosts over a decade of regional circulation, suggesting that its broad host range is a stable and intrinsic trait rather than a recent adaptation (23, 24). In addition to the plasmid factors contributing to its success, there are likely environmental factors. Continued, and sometimes inappropriate, use of carbapenem antibiotics within hospitals applies consistent selective pressure, further amplifying KPC-producers (18, 25). Additionally, lapses in infection control and patient movement between hospitals and long-term care facilities have repeatedly enabled clonal and plasmid-level spread (18). Together, these factors underscore why KPC-positive plasmids can maintain prevalence in a region for many years despite infection control efforts.

Investigating the prevalent plasmids in the region, provides insights into the plasmid epidemiology among various species. Though multiple tools exist for plasmid classification within Enterobacterales, they differ in methodology. PlasmidFinder, for example, classifies plasmids based on incompatibility (Inc) types, whereas MOB-suite uses mash distance-based clustering to group similar plasmids (26, 27). PlasX classifies plasmids into plasmid taxonomic units (PTUs) using average nucleotide identity (ANI), and MobMess relies on mobility gene content and sequence identity to assign plasmid mobility classes and perform pairwise comparisons(28). However, the accuracy of these classifications is highly dependent on the underlying databases used. Tools with large, diverse, and well-represented plasmid reference datasets tend to provide more precise and meaningful clustering of related plasmids(15, 16). In the present study, MOB-suite was used to classify plasmids into clusters to define the overall distribution of plasmids within each species. Further, we integrated complete plasmid sequences into the MOB-suite database to enhance classification and improve clustering by increasing the plasmid representation in the database. This addition resulted in 71% of plasmids mapping to a reference within the study collection and 50% of the plasmids mapping with a mash distance of <0.005, while with the standard reference database only 30% of plasmids mapped with a mash distance <0.005. This finding supports the potential to use targeted long-read sequencing to enhance tracking of locally circulating plasmids.

Lastly, we believe that our observation for the key role of an epidemic plasmid, harbored by diverse strains and species, has implications on both infection prevention and surveillance activities. First, the role of uncommon organisms in plasmid transfer networks highlights the need to track not just multidrug-resistant organisms, but to also consider organisms with more limited resistance profiles. Tracking these atypical species is essential to understanding the scope of plasmid-mediated outbreaks, as well as for making epidemiological connections among non-clonally linked cases, and thereby enabling tracking the pathways of spread. Moreover, given their typical low pathogenic nature, we are likely underestimating their prevalence, and in turn their potential contribution to plasmid dissemination into genetic backgrounds with potential for clonal spread. In the present study, detection of the AK975 plasmid among species such as *C. werkmanii*, *C. europaeus*, and *K. grimontii* was only possible through genomic surveillance. Importantly, since AK975 was absent among major clones like *K. pneumoniae* ST258 and *E. coli* ST131, surveillance of these common species will not significantly contribute to tracking the spread of this *bla*_KPC_ plasmid in Michigan. Thus, identifying the genomic mechanisms underlying resistance spread is crucial for effective intervention and containment. Second, the key role of plasmid transfer and organisms not typically causing healthcare infections, points to the potential need for distinct intervention strategies. Previous studies of plasmid outbreaks at individual healthcare institutions have suggested a potential role for hospital plumbing and sinks, in both creating a favorable environment for plasmid transfer and acting as reservoirs for subsequent transmission (2, 11, 13, 29). Whole-genome sequencing at the University of Virginia Health Systems detected KPC plasmid outbreaks in uncommon species from sink traps (30). Thus, the presence of epidemic plasmids, such as the AK975 plasmid observed in Michigan, may warrant increased attention to environmental surveillance and disinfection when contemplating interventions to reduce the burden of resistance. Infection control strategies in these scenarios must include eliminating environmental reservoirs and implementing interventions beyond patient-centered approaches (29).

## Conclusion

This study offers valuable insights into the relative roles of clonal spread and plasmid transfer in maintaining the endemicity of blaKPC in a statewide healthcare network over a decade. We observe a key role of epidemic lineages, contributing to KPC dissemination through clonal amplification of diverse plasmids, including the endemic AK975 plasmid. In parallel to clonal spread of epidemic lineages, we observe a multi-species plasmid-mediated epidemic, with the AK975 plasmid being found across both epidemic and non-epidemic lineages in 20 species. This broad distribution underscores the importance of surveillance efforts that extend beyond common pathogens, and track not just lineages with epidemic potential, but also mobile elements with epidemic potential. Our findings support the integration of plasmid surveillance into regional epidemiological studies, especially when plasmids with ARGs show inter-species spread. Future studies are needed to improve our understanding of the defining features of epidemic plasmids, the nature of the organism reservoirs that maintain them and the inter-organisms transfer patterns that mediate their broad spread.

## Materials and methods

### Bacterial Isolates

KPC-producing isolates were submitted by the HCFs to the Bureau of Laboratories at the Michigan Department of Health and Human Services starting in 2013. From 2013- 2021, healthcare facility submission of carbapenem resistant Enterobacterales (CRE) isolates to the state public health laboratory was voluntary, while from 2022 onwards submission of CRE isolates was required. In the current study we performed whole-genome sequencing on all isolates submitted between 2013 and 2022. The isolates were obtained from various clinical specimens including but not limited to blood, urine, sputum, rectal swabs and stool. Organism identifications were confirmed by MALDI-TOF, carbapenem resistance confirmed using Sensititre™ Gram Negative GNX2F AST Plate (Thermofisher), and carbapenemase production determined using the modified carbapenem inactivation method (mCIM)(29) and CDC-developed PCR assays (https://www.cdc.gov/gram-negative-bacteria/php/laboratories/index.html) .

### Genome sequencing and assembly

For Illumina sequencing, all 1082 KPC-producing bacteria had DNA extractions performed on Epimotion liquid handling robots using the QIAGEN MagAttract microbial DNA kit. Genomic libraries were prepared with the QIASeq FX DNA library prep kit and sequenced at the University of Michigan Advanced Genomics Core on an Illumina NovaSeq 6000, with 150-bp paired-end reads. The short-reads were then analyzed with the Quality Control and Contamination Detection (QCD) workflow as described in https://github.com/Snitkin-Lab-Umich/QCD. Briefly, raw sequencing reads were trimmed using Trimmomatic v.0.39 to remove adapters and low-quality bases (31). Trimmed high-quality reads were assembled using Spades v.3.15.3 (32) and annotated using Prokka v.1.14.5 (33). Criteria for passing QC included sequencing depth greater than 50X and number of contigs less than 500.

Among the 1065 genomes that passed quality assessment for short-read sequencing, a subset of 599 isolates were chosen for long-read sequencing that represented unique combinations of species, year of isolation, *bla*_KPC_ allele, plasmid cluster (described below) and the Tn*4401* variant associated with *bla*_KPC_ (described below). The isolates chosen for long-read sequencing included those that differed by >5 SNPs in their core-genome. For ONT sequencing, DNA extraction was performed using the Promega WizardⓇ DNA extraction kit (Promega, USA) as per manufacturer’s instructions and sequenced by Plasmidsaurus on a PromethION sequencer using ONT’s rapid barcoding kit V14. The Nanopore reads were subjected to nanoQC for quality control and assembled using Flye (v2.9.5) (https://github.com/Snitkin-Lab-Umich/nanoQC) (34). Polished assemblies combining short and long reads were generated using the nanosake pipeline (https://github.com/Snitkin-Lab-Umich/Nanosake). Briefly, polished assemblies were generated using long reads assembled by Flye (v2.9.5), which is polished using Medaka (v1.2.0, https://github.com/nanoporetech/medaka) and combining this with clean trimmed illumina reads using Polypolish (v0.6.0) to generate final assemblies (34–36).

### Whole genome analysis

Species were identified using Skani (v0.2.1) (https://github.com/bluenote-1577/skani). In the present study, *Enterobacter cloacae* species complex includes the species *E. cloacae*, *E. hormaechei, E. kobei* and *E. asburiae*. *Citrobacter freundii* species complex includes the species *C. freundii*, *C. brakii* and *C. koseri*. The presence of antimicrobial resistance genes, plasmids and transposons were analyzed using AMRFinder Plus (v3.11.4), MOB-suite (v3.1.17) and TETyper (v1.1) respectively(26, 37, 38). Multi-locus Sequence Typing (MLST, 2.23.0) was performed as per MLST (39). The primary plasmid clusters defined by MOB-suite from the short-read assemblies, were then considered to select the isolates for long-read sequencing as mentioned above.

The core genome phylogeny of epidemic lineages such as *K. pneumoniae* ST11 and ST258, *E. coli* ST131, *C. freundii* ST22 and *E. cloacae* complex ST78 and ST171 were generated using SNPkit (https://github.com/Snitkin-Lab-Umich/snpkit-smk). Briefly, trimmed sequencing reads were mapped to reference genomes using the Burrows– Wheeler Aligner-MEM v.0.7.17 (40) and variants were called and filtered using Samtools v.1.11(41). Variants were filtered from raw results using GATK’s VariantFiltration (QUAL, >100; MQ, >50; >=10 reads supporting variant; and FQ, <0.025). In addition, a custom python script was used to filter out single- nucleotide variants that were: (i) <5 base pairs (bp) in proximity to indels that were identified by GATK HaplotypeCaller, (ii) in a phage region identified by Phastest (42) or (iii) they resided in tandem repeats of length greater than 20 bp as determined using the exact-tandem program in MUMmer(43). The reference genomes used for variant calling include <u>CP066523.1</u> for *K. pneumoniae* ST11, <u>NZ_CP008827.1</u> for *K. pneumoniae* ST258, <u>GCA_900448475.1</u> for *E. coli* ST131 and <u>CP012165.1</u> for *E. cloacae* ST171. For *C. freundii* ST22 *E. hormaechei* ST78, internal references from the study collection were included such as MI_KPC_1033 (SRS26152792) and MI_KPC_249 (SRS26152745) respectively (earliest isolate of respective ST from the study). Cognac was used for constructing phylogenies of non-epidemic lineages of major species(44).

### Plasmid analysis

Initially, the short-read assemblies were run using the standard MOB-suite (v3.1.17) database and the primary cluster was used to choose isolates for long-read as mentioned above. Once the hybrid assemblies were generated, the MOB-suite database was updated to include all circular plasmids from the study collection in order to improve the plasmid clustering and create a database that included the KPC plasmids from Michigan. MOB-suite was re-run for the short reads using the updated MOB-suite and the clusters were redefined. The secondary clusters that were defined by MOB-suite are referred to in the manuscript. SNP based phylogeny was constructed for the major KPC plasmid cluster, AK975, using Parsnp (45). The reference plasmid used for this was CP011598 which is the nearest reference in the MOB-suite database. All the complete circular KPC plasmids (n=384) obtained from hybrid assemblies were used to construct a mash-distance based phylogeny (46).

## Data analysis and visualization

The analyses and data visualization were performed with R studio using the packages ggplot2, tidyverse, lubridate, ggtree, phytools, ape and readxl.

## Data availability

Whole genome sequences were deposited under BioProject accession number PRJNA1305037.

## Acknowledgements

Funding was provided as part of the Michigan Sequencing and Academic Partnerships for Public Health Innovation and Response (MI-SAPPIRE) initiative at the Michigan Department of Health and Human Services (MDHHS) which is supported with funds from the Centers for Disease Control and Prevention through the Epidemiology and Laboratory Capacity for Prevention and Control of Emerging Infectious Disease Enhancing Detection Expansion (6NU50CK000510-02-07). ESS was also supported by the National Institutes of Health grant 1U19AI181767.

## References

1. Yigit H, Queenan AM, Anderson GJ, Domenech-Sanchez A, Biddle JW, Steward CD, Alberti S, Bush K, Tenover FC. 2001. Novel Carbapenem-Hydrolyzing β-Lactamase, KPC- 1, from a Carbapenem-Resistant Strain of Klebsiella pneumoniae. Antimicrob Agents Chemother 45:1151–1161.

2. Falagas ME, Tansarli GS, Karageorgopoulos DE, Vardakas KZ. 2014. Deaths Attributable to Carbapenem-Resistant Enterobacteriaceae Infections. Emerg Infect Dis 20:1170–1175.

3. Qureshi ZA, Paterson DL, Potoski BA, Kilayko MC, Sandovsky G, Sordillo E, Polsky B, Adams-Haduch JM, Doi Y. 2012. Treatment outcome of bacteremia due to KPC-producing Klebsiella pneumoniae: superiority of combination antimicrobial regimens. Antimicrob Agents Chemother 56:2108–2113.

4. Bertagnolio S, Dobreva Z, Centner CM, Olaru ID, Donà D, Burzo S, Huttner BD, Chaillon A, Gebreselassie N, Wi T, Hasso-Agopsowicz M, Allegranzi B, Sati H, Ivanovska V, Kothari KU, Balkhy HH, Cassini A, Hamers RL, Weezenbeek KV, Aanensen D, Alanio A, Alastruey-Izquierdo A, Alemayehu T, Al-Hasan M, Allegaert K, Al-Maani AS, Al-Salman J, Alshukairi AN, Amir A, Applegate T, Araj GF, Villalobos MA, Årdal C, Ashiru-Oredope D, Ashley EA, Babin F-X, Bachmann LH, Bachmann T, Baker KS, Balasegaram M, Bamford C, Baquero F, Barcelona LI, Bassat Q, Bassetti M, Basu S, Beardsley J, Vásquez GB, Berkley JA, Bhatnagar AK, Bielicki J, Bines J, Bongomin F, Bonomo RA, Bradley JS, Bradshaw C, Brett A, Brink A, Brown C, Brown J, Buising K, Carson C, Carvalho AC, Castagnola E, Cavaleri M, Cecchini M, Chabala C, Chaisson RE, Chakrabarti A, Chandler C, Chandy SJ, Charani E, Chen L, Chiara F, Chowdhary A, Chua A, Chuki P, Chun DR, Churchyard G, Cirillo D, Clack L, Coffin SE, Cohn J, Cole M, Conly J, Cooper B, Corso A, Cosgrove SE, Cox H, Daley CL, Darboe S, Darton T, Davies G, Egea V de, Dedeić-Ljubović A, Deeves M, Denkinger C, Dillon J-AR, Dramowski A, Eley B, Esposito SMR, Essack SY, Farida H, Farooqi J, Feasey N, Ferreyra C, Fifer H, Finlayson H, Frick M, Gales AC, Galli L, Gandra S, Gerber JS, Giske C, Gordon B, Govender N, Guessennd N, Guindo I, Gurbanova E, Gwee A, Hagen F, Harbarth S, Haze J, Heim J, Hendriksen R, Heyderman RS, Holt KE, Hönigl M, Hook EW, Hope W, Hopkins H, Hughes G, Ismail G, Issack MI, Jacobs J, Jasovský D, Jehan F, Pearson AJ, Jones M, Joshi MP, Kapil A, Kariuki S, Karkey A, Kearns GL, Keddy KH, Khanna N, Kitamura A, Kolho K-L, Kontoyiannis DP, Kotwani A, Kozlov RS, Kranzer K, Kularatne R, Lahra MM, Langford BJ, Laniado-Laborin R, Larsson J, Lass-Flörl C, Doare KL, Lee H, Lessa F, Levin AS, Limmathurotsakul D, Lincopan N, Vecchio AL, Lodha R, Loeb M, Longtin Y, Lye DC, Mahmud AM, Manaia C, Manderson L, Mareković I, Marimuthu K, Martin I, Mashe T, Mei Z, Meis JF, Melo FALTD, Mendelson M, Miranda AE, Moore D, Morel C, Moremi N, Moro ML, Moussy F, Mshana S, Mueller A, Ndow FJ, Nicol M, Nunn A, Obaro S, Obiero CW, Okeke IN, Okomo U, Okwor TJ, Oladele R, Omulo S, Ondoa P, Canese JMO de, Ostrosky-Zeichner L, Padoveze MC, Pai M, Park B, Parkhill J, Parry CM, Peeling R, Peixe LMSV, Perovic O, Pettigrew MM, Principi N, Pulcini C, Puspandari N, Rawson T, Reddy DL, Reddy K, Redner P, Tudela JLR, Rodríguez-Baño J, Katwyk SRV, Roilides E, Rollier C, Rollock L, Ronat J-B, Ruppe E, Sadarangani M, Salisbury D, Salou M, Samison LH, Sanguinetti M, Sartelli M, Schellack N, Schouten J, Schwaber MJ, Seni J, Senok A, Shafer WM, Shakoor S, Sheppard D, Shin J-H, Sia S, Sievert D, Singh I, Singla R, Skov RL, Soge OO, Sprute R, Srinivasan A, Srinivasan S, Sundsfjord A, Tacconelli E, Tahseen S, Tangcharoensathien V, Tängdén T, Thursky K, Thwaites G, Peral RT de S, Tong D, Tootla HD, Tsioutis C, Turner KM, Turner P, Omar SV, Sande WW van de, Hof S van den, Doorn R van, Veeraraghavan B, Verweij P, Wahyuningsih R, Wang H, Warris A, Weinstock H, Wesangula E, Whiley D, White PJ, Williams P, Xiao Y, Moscoso MY, Yang HL, Yoshida S, Yu Y, Żabicka D, Zignol M. 2024. WHO global research priorities for antimicrobial resistance in human health. Lancet Microbe 0.

5. Wilhelm CM, Antochevis LC, Magagnin CM, Arns B, Vieceli T, Pereira DC, Lutz L, de Souza ÂC, dos Santos JN, Guerra RR, Medeiros GS, Santoro L, Falci DR, Rigatto MH, Barth AL, Martins AF, Zavascki AP. 2024. Susceptibility evaluation of novel beta- lactam/beta-lactamase inhibitor combinations against carbapenem-resistant *Klebsiella pneumoniae* from bloodstream infections in hospitalized patients in Brazil. J Glob Antimicrob Resist 38:247–251.

6. Camargo CH, Yamada AY, de Souza AR, Cunha MPV, Ferraro PSP, Sacchi CT, dos Santos MB, Campos KR, Tiba-Casas MR, Freire MP, Barretti P. 2023. Genomic analysis and antimicrobial activity of β-lactam/β-lactamase inhibitors and other agents against KPC- producing Klebsiella pneumoniae clinical isolates from Brazilian hospitals. Sci Rep 13:14603.

7. Marsh JW, Mustapha MM, Griffith MP, Evans DR, Ezeonwuka C, Pasculle AW, Shutt KA, Sundermann A, Ayres AM, Shields RK, Babiker A, Cooper VS, Van Tyne D, Harrison LH. 2019. Evolution of Outbreak-Causing Carbapenem-Resistant Klebsiella pneumoniae ST258 at a Tertiary Care Hospital over 8 Years. mBio 10:10.1128/mbio.01945-19.

8. Kitchel B, Rasheed JK, Patel JB, Srinivasan A, Navon-Venezia S, Carmeli Y, Brolund A, Giske CG. 2009. Molecular Epidemiology of KPC-Producing Klebsiella pneumoniae Isolates in the United States: Clonal Expansion of Multilocus Sequence Type 258. Antimicrob Agents Chemother 53:3365–3370.

9. Gomez-Simmonds A, Hu Y, Sullivan SB, Wang Z, Whittier S, Uhlemann A-C. 2016. Evidence from a New York City hospital of rising incidence of genetically diverse carbapenem-resistant Enterobacter cloacae and dominance of ST171, 2007–14. J Antimicrob Chemother 71:2351–2353.

10. Gomez-Simmonds A, Annavajhala MK, Wang Z, Macesic N, Hu Y, Giddins MJ, O’Malley A, Toussaint NC, Whittier S, Torres VJ, Uhlemann A-C. 2018. Genomic and Geographic Context for the Evolution of High-Risk Carbapenem-Resistant Enterobacter cloacae Complex Clones ST171 and ST78. mBio 9:e00542–18.

11. Salamzade R, Manson AL, Walker BJ, Brennan-Krohn T, Worby CJ, Ma P, He LL, Shea TP, Qu J, Chapman SB, Howe W, Young SK, Wurster JI, Delaney ML, Kanjilal S, Onderdonk AB, Bittencourt CE, Gussin GM, Kim D, Peterson EM, Ferraro MJ, Hooper DC, Shenoy ES, Cuomo CA, Cosimi LA, Huang SS, Kirby JE, Pierce VM, Bhattacharyya RP, Earl AM. 2022. Inter-species geographic signatures for tracing horizontal gene transfer and long-term persistence of carbapenem resistance. Genome Med 14:37.

12. Sugita K, Aoki K, Komori K, Nagasawa T, Ishii Y, Iwata S, Tateda K. 2021. Molecular Analysis of blaKPC-2-Harboring Plasmids: Tn4401a Interplasmid Transposition and Tn4401a-Carrying ColRNAI Plasmid Mobilization from Klebsiella pneumoniae to Citrobacter europaeus and Morganella morganii in a Single Patient. mSphere 6:e00850–21.

13. Brandt C, Viehweger A, Singh A, Pletz MW, Wibberg D, Kalinowski J, Lerch S, Müller B, Makarewicz O. 2019. Assessing genetic diversity and similarity of 435 KPC-carrying plasmids. Sci Rep 9:N.PAG-N.PAG.

14. Wick RR, Holt KE. 2021. Benchmarking of long-read assemblers for prokaryote whole genome sequencing. F1000Research 8:2138.

15. Gomez-Simmonds A, Annavajhala MK, Seeram D, Hokunson TW, Park H, Uhlemann A-C. 2024. Genomic epidemiology of carbapenem-resistant Enterobacterales at a New York City hospital over a ten-year period reveals complex plasmid-clone dynamics and evidence for frequent horizontal transfer of *bla* _KPC_. Genome Res gr.279355.124.

16. Lerminiaux N, Mitchell R, Bartoszko J, Davis I, Ellis C, Fakharuddin K, Hota SS, Katz K, Kibsey P, Leis JA, Longtin Y, McGeer A, Minion J, Mulvey M, Musto S, Rajda E, Smith SW, Srigley JA, Suh KN, Thampi N, Tomlinson J, Wong T,Mataseje L. Plasmid genomic epidemiology of blaKPC carbapenemase-producing Enterobacterales in Canada, 2010– 2021. Antimicrob Agents Chemother 67:e00860–23.

17. Mathers AJ, Peirano G, Pitout JDD. 2015. The Role of Epidemic Resistance Plasmids and International High-Risk Clones in the Spread of Multidrug-Resistant Enterobacteriaceae. Clin Microbiol Rev 28:565–591.

18. Munoz-Price LS, Quinn JP. 2009. The spread of Klebsiella pneumoniae carbapenemases: a tale of strains, plasmids, and transposons. Clin Infect Dis Off Publ Infect Dis Soc Am 49:1739–1741.

19. San Millan A, MacLean RC. 2017. Fitness Costs of Plasmids: a Limit to Plasmid Transmission. Microbiol Spectr 5:10.1128/microbiolspec.mtbp-0016–2017.

20. Harrison E, Guymer D, Spiers AJ, Paterson S, Brockhurst MA. 2015. Parallel Compensatory Evolution Stabilizes Plasmids across the Parasitism-Mutualism Continuum. Curr Biol 25:2034–2039.

21. Porse A, Schønning K, Munck C, Sommer MOA. 2016. Survival and Evolution of a Large Multidrug Resistance Plasmid in New Clinical Bacterial Hosts. Mol Biol Evol 33:2860– 2873.

22. Vogwill T, MacLean RC. 2015. The genetic basis of the fitness costs of antimicrobial resistance: a meta-analysis approach. Evol Appl 8:284–295.

23. San Millan A. 2018. Evolution of Plasmid-Mediated Antibiotic Resistance in the Clinical Context. Trends Microbiol 26:978–985.

24. Hall JPJ, Brockhurst MA, Harrison E. 2017. Sampling the mobile gene pool: innovation via horizontal gene transfer in bacteria. Philos Trans R Soc Lond B Biol Sci 372:20160424.

25. van Duin D, Doi Y. 2016. The global epidemiology of carbapenemase-producing Enterobacteriaceae. Virulence 8:460–469.

26. Robertson J, Nash JHE. 2018. MOB-suite: software tools for clustering, reconstruction and typing of plasmids from draft assemblies. Microb Genomics 4:e000206.

27. Carattoli A, Hasman H. 2020. PlasmidFinder and In Silico pMLST: Identification and Typing of Plasmid Replicons in Whole-Genome Sequencing (WGS). Methods Mol Biol Clifton NJ 2075:285–294.

28. Yu MK, Fogarty EC, Eren AM. 2024. Diverse plasmid systems and their ecology across human gut metagenomes revealed by PlasX and MobMess. Nat Microbiol 9:830–847.

29. Tsukada M, Miyazaki T, Aoki K, Yoshizawa S, Kondo Y, Sawa T, Murakami H, Sato E, Tomida M, Otani M, Kumade E, Takamori E, Kambe M, Ishii Y, Tateda K. 2024. The outbreak of multispecies carbapenemase-producing Enterobacterales associated with pediatric ward sinks: IncM1 plasmids act as vehicles for cross-species transmission. Am J Infect Control S0196655324001019.

30. Mathers AJ, Li TJX, He Q, Narendra S, Stoesser N, Eyre DW, Walker AS, Barry KE, Castañeda-Barba S, Huang FW, Parikh H, Kotay S, Crook DW, Reidys C. 2024. Developing a framework for tracking antimicrobial resistance gene movement in a persistent environmental reservoir. Npj Antimicrob Resist 2:1–12.

31. Bolger AM, Lohse M, Usadel B. 2014. Trimmomatic: a flexible trimmer for Illumina sequence data. Bioinformatics 30:2114–2120.

32. Prjibelski A, Antipov D, Meleshko D, Lapidus A, Korobeynikov A. 2020. Using SPAdes De Novo Assembler. Curr Protoc Bioinforma 70:e102.

33. Seemann T. 2014. Prokka: rapid prokaryotic genome annotation. Bioinformatics 30:2068– 2069.

34. Kolmogorov M, Bickhart DM, Behsaz B, Gurevich A, Rayko M, Shin SB, Kuhn K, Yuan J, Polevikov E, Smith TPL, Pevzner PA. 2020. metaFlye: scalable long-read metagenome assembly using repeat graphs. Nat Methods 17:1103–1110.

35. Wick RR, Holt KE. 2022. Polypolish: Short-read polishing of long-read bacterial genome assemblies. PLOS Comput Biol 18:e1009802.

36. Prior K, Becker K, Brandt C, Cabal Rosel A, Dabernig-Heinz J, Kohler C, Lohde M, Ruppitsch W, Schuler F, Wagner GE, Mellmann A. 2025. Accurate and reproducible whole-genome genotyping for bacterial genomic surveillance with Nanopore sequencing data. J Clin Microbiol 63:e0036925.

37. Feldgarden M, Brover V, Gonzalez-Escalona N, Frye JG, Haendiges J, Haft DH, Hoffmann M, Pettengill JB, Prasad AB, Tillman GE, Tyson GH, Klimke W. 2021. AMRFinderPlus and the Reference Gene Catalog facilitate examination of the genomic links among antimicrobial resistance, stress response, and virulence. 1. Sci Rep 11:12728.

38. Sheppard AE, Stoesser N, German-Mesner I, Vegesana K, Sarah Walker A, Crook DW, Mathers AJ. 2018. TETyper: a bioinformatic pipeline for classifying variation and genetic contexts of transposable elements from short-read whole-genome sequencing data. Microb Genomics 4:e000232.

39. Jolley KA, Maiden MC. 2010. BIGSdb: Scalable analysis of bacterial genome variation at the population level. BMC Bioinformatics 11:595.

40. Li H, Durbin R. 2009. Fast and accurate short read alignment with Burrows–Wheeler transform. Bioinformatics 25:1754–1760.

41. Danecek P, Bonfield JK, Liddle J, Marshall J, Ohan V, Pollard MO, Whitwham A, Keane T, McCarthy SA, Davies RM, Li H. 2021. Twelve years of SAMtools and BCFtools. GigaScience 10:giab008.

42. Arndt D, Grant JR, Marcu A, Sajed T, Pon A, Liang Y, Wishart DS. 2016. PHASTER: a better, faster version of the PHAST phage search tool. Nucleic Acids Res 44:W16–W21.

43. Marçais G, Delcher AL, Phillippy AM, Coston R, Salzberg SL, Zimin A. 2018. MUMmer4: A fast and versatile genome alignment system. PLoS Comput Biol 14:e1005944.

44. Crawford RD, Snitkin ES. 2021. cognac: rapid generation of concatenated gene alignments for phylogenetic inference from large, bacterial whole genome sequencing datasets. BMC Bioinformatics 22:70.

45. Kille B, Nute MG, Huang V, Kim E, Phillippy AM, Treangen TJ. 2024. Parsnp 2.0: scalable core-genome alignment for massive microbial datasets. Bioinformatics 40:btae311.

46. Katz L, Griswold T, Morrison S, Caravas J, Zhang S, Bakker H, Deng X, Carleton H. 2019. Mashtree: a rapid comparison of whole genome sequence files. J Open Source Softw 4:1762.

